# The Missing Memory Imprint

**DOI:** 10.1101/2025.05.05.652252

**Authors:** Howard Bowman, Arthur Assant, Alberto Aviles

## Abstract

We provide evidence that the brain searches for salient stimuli below the level of conscious awareness. We show that, while targets are found very efficiently in Rapid Serial Visual Presentation (RSVP), distractors do not leave a strong memory trace, especially if the memory probe is unexpected. This fits with traditional theories of late attentional selection, whereby “correct rejections” of non-targets are performed with very little processing cost. Our findings are also consistent with the tokenized percept theory of conscious perception, one element of which is that conscious awareness is required to provide sustained representations.

## Introduction

Bowman et al (2013) introduced the term *subliminal salience search* (SSS) to describe the capacity that we have to detect and identify salient stimuli in our environments, with perception of salient stimuli in Rapid Serial Visual Presentation (RSVP) streams used as a representative case. Clearly, the loaded term here is *subliminal*: do we definitively know that the brain is finding salient stimuli in RSVP streams below the threshold of awareness? Indeed, it could be that the term *liminal* salience search is more appropriate, i.e. that percepts are fleetingly experienced in RSVP, but are not strictly subconscious. The objective of this paper is to provide evidence that the sub- in subliminal can be appropriately used.

The perceptual experience associated with viewing RSVP is certainly strange, with a jumble of stimuli appearing rapidly, with vividness of percept varying according to changes in masking strength between items. However, objectively measured performance at seeing salient stimuli, across a range of possible dimensions of salience, is excellent. A classic example of such high performance is Potter et al, 2014 (see figure 1) where participants searched RSVP streams of pictures on the basis of meaning with stimuli presented at 78 per second; other examples include (Potter, 1976; Barnard et al, 2004; Most et al, 2005; Bowman et al, 2013), and many others.

**Figure 1:**
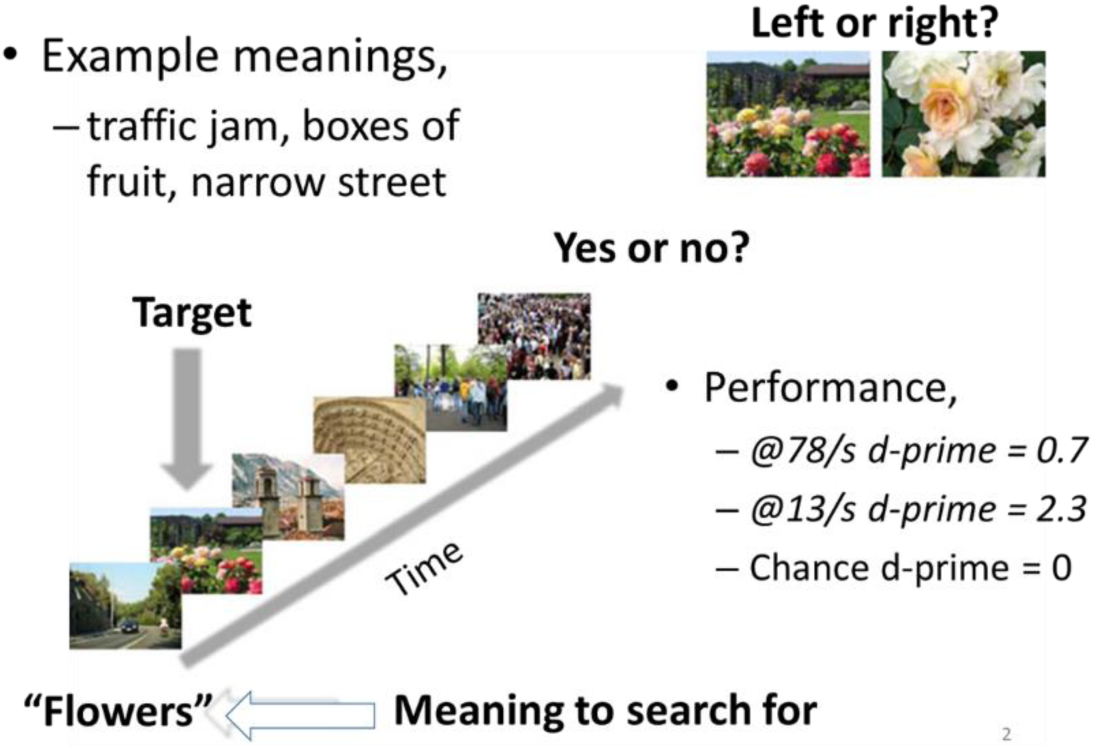
Illustration of brain’s impressive capacity to search RSVP streams. Participants were given a meaning to search for in an RSVP stream of pictures. At stream-end, they performed a presence/ absence judgement on the occurrence of a picture consistent with the target meaning and then a 2-alternative forced choice on its identity. Notably, even when pictures were presented at 78 per second, detection performance was well above chance, with a d-prime of 0.7. Figure reprinted from Potter, M. C., Wyble, B., Hagmann, C. E., & McCourt, E. S. (2014). Detecting meaning in RSVP at 13 ms per picture. Attention, Perception, & Psychophysics, 76(2), 270-279.

Thus, it is clear that the human brain is very good at finding salient (e.g. target) stimuli in RSVP streams. In this sense, the brain is performing a search when viewing such streams, i.e. it is seeking out stimuli that are of interest to it. When such a stimulus is found, it is more fully perceived and then enters into working memory, enabling it to be reported.

The key logic underlying the approach taken in this paper is that if the process of finding salient stimuli in RSVP is conscious, the case of *liminal* salience search, memory imprints should be left for many of the *not* salient stimuli presented in the stream. Such a finding would be consistent with *late selection* theories of attentional processes (Deutsch & Deutsch, 1963; Duncan, 1980). However, the role of conscious experience is more overtly emphasized in our current work; that is, that the late selection of classic *attentional* theories precedes and initiates a *conscious* perceptual event. We will return to the relationship to late selection in the Discussion.

In terms of RSVP experiments, the late selection/ *sub*liminal salience search perspective suggests interest in the memory traces left by arbitrary distractors. In particular, we can follow the logic that we introduced in Bowman & Avilés (2021), which proceeds as follows. Since the position at which targets can appear in streams is randomly varied across trials, a high detection performance, where, say, 90% of trials are correctly responded to (hits or correct rejects), would require a very high percentage of the items presented in the stream to be processed sufficiently to determine if they are targets. If they are judged to be targets, they are encoded into working memory, but what happens to stimuli that are not judged to be targets? Under the liminal salience search hypothesis, some sort of memory trace would be left for a very large number of items, even if only one of these was a target.

The question of the memory trace left in RSVP was first investigated in Mary Potter’s classic work in the 70’s (Potter, 1976). More recently, we showed that free recall was very poor for RSVP stream distractors (Bowman & Avilés, 2021). However, neither of these previous studies probed memory in a fashion that could truly extrapolate to an arbitrary distractor in an RSVP stream, which is what is of interest for the Subliminal Salience Search (SSS) hypothesis. Thus, the central issue remains moot. This is what we seek to resolve in this article.

In the studies reported here, we probe recognition and detection, two yes/no decision tasks, enabling us to compare d-primes for detection to those for recognition. However, this does not resolve the key experimental difficulty, which is that the memory traces left could be very brief, i.e. the percept created by a distractor could be forgotten very rapidly. That is, strictly speaking, there was an imprint generated, but it evaporated long before being probed for at the end of the RSVP stream.

We resolve this issue by, firstly, randomly varying the length of RSVP streams, in order that participants cannot build up an expectation for when the end of stream would come, and, secondly, by just probing memory for the last distractor presented. This gives the shortest time for forgetting before probing memory. Although, of course, this distractor has to be masked.

We explore two Stimulus Onset Asynchronies (SOAs), 117ms and 350ms. The first of these is a typical RSVP rate^1^, with stimuli presented “on the fringe of awareness”, or, indeed, below it, while the second should be a conscious regime, in which the vast majority of stimuli would generate a conscious experience.

Our first experiment compares recognition to detection for a range of (last item) masks – words, pseudo-words and random symbol strings – as well as a “boundary” condition in which no mask was presented. This enables us to determine whether reduced recognition performance compared to detection was robust across types of mask and additionally whether masking was more “perceptually-determined” (as opposed to conceptually) at short, rather than at long SOAs.

Our second experiment seeks to test for memory imprints, when the brain is in its “native” (sub)liminal search state. Specifically, we are interested in the memory traces left by distractors, when the brain is solely engaged in search for target stimuli, i.e. is just performing the detection task. Whether instructed to do so or not, as soon as memory has been probed once during an experiment, participants will be attempting to encode distractors, and they will effectively be performing a dual task: “detect targets *and* encode distractors”. To resolve this issue, we use the approach introduced in Chen and Wyble (2015) and introduce a surprise recognition memory probe in a single trial during a pure detection experiment.

## Experiment 1

### Methods

#### Participants

30 (mean age=19.33, sd=1.24) undergraduate students (8 males) of the University of Birmingham took part in the experiment in exchange for course credits. All were native English speakers and had normal or corrected-to-normal vision. The experiment conformed to British Psychological Society criteria for the ethical conduct of research and ethical procedures of the School of Psychology at the University of Birmingham. The experiments performed here were approved by the Science, Technology, Engineering and Mathematics Ethical Review Committee at the University of Birmingham, under ethics programme ERN_20-1291P.

### Materials

1000 English words were selected from (Warriner, Kuperman, & Brysbaert, 2013) to serve as distractors. 224 from the same source were selected as Targets for the detection task. 224 words were selected for the recognition test (half served as Old Words and half as New Words). Four lists of stimuli were created so that the Old Words were presented in each of the masking conditions. Additionally, we generated 56 pseudo words by changing two letters of each of a new set of 56 words.

### Procedure

We presented RSVP streams on a 24’’ LCD screen (refresh rate: 60Hz, resolution: 1920 x 1080) using custom Psychopy3 (Peirce et al., 2019) scripts running under Python 3.6. Stimuli were Arial white characters of 1.3^°^ of visual angle in height on a grey background (40% white). Participants were seated at 60cm from the screen. The experiment was conducted individually in a quiet room. Each experimental session consisted of 224 RSVP trials.

The format of the experiment is shown in figure 2. Each trial started with the presentation of the instruction “Search for the word:” in the upper part of the screen along with the target word presented at the centre of the screen. Participants were instructed to press the “space bar” after reading the target word to continue. The starting item was a fixation cross (+) presented for 500ms at the centre of the screen. Then a sequence of 9 to 19 (varied uniformly randomly) words was presented at an SOA of 117ms or 350ms. The finishing item, presented for the same SOA as the other stream items, varied depending on the masking condition:

**Figure 2:**
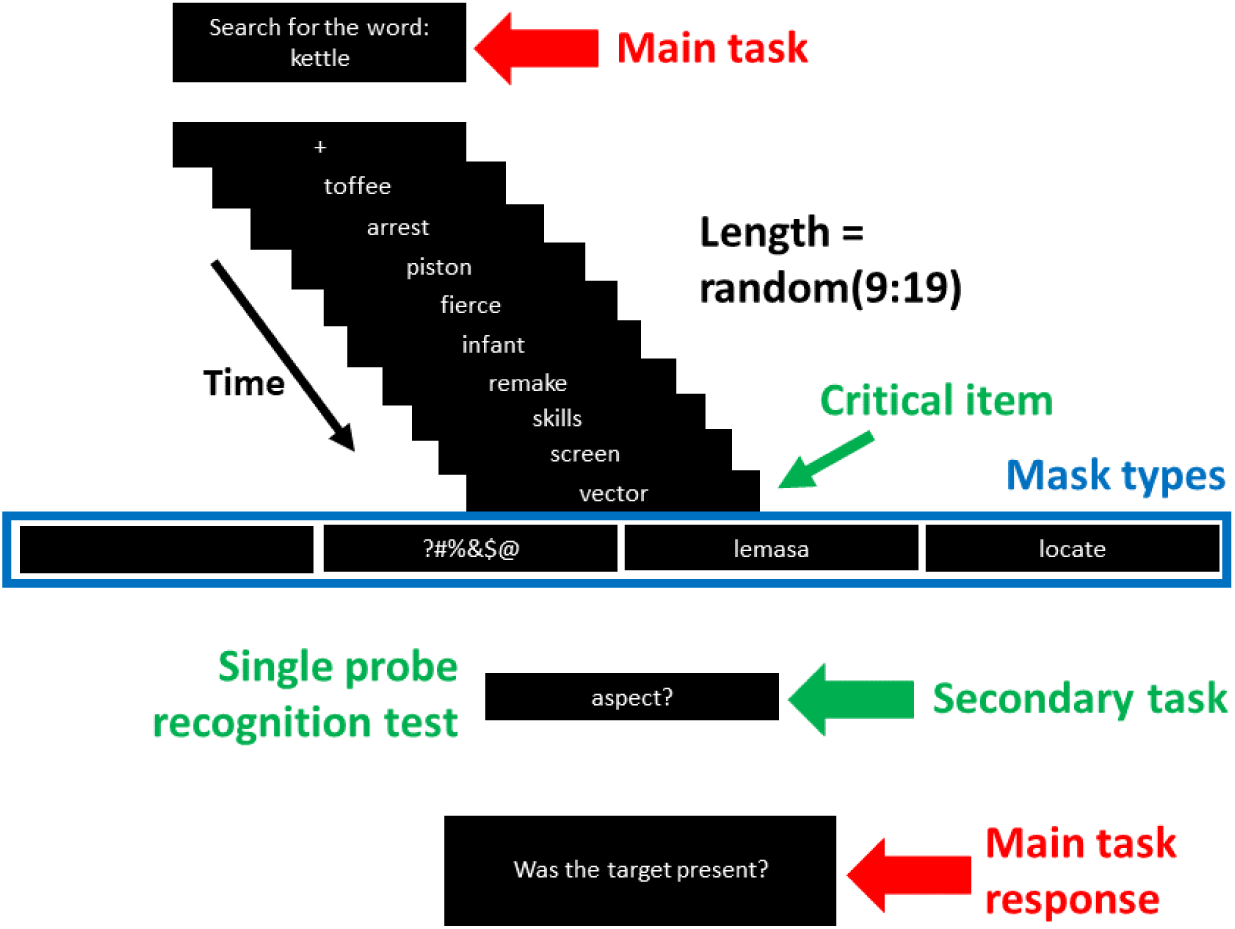
experiment 1 procedure: RSVP streams of words were presented, terminated by a mask. Streams were stopped randomly from the 9th to the 19th item. The main task was to detect the specified word and perform a presence/ absence judgement on it at stream end (see Main task response). Additionally, in a Secondary task, participants performed a single-probe recognition test on the critical item: the last stream item before the mask. Four different mask types were used.

Word: a random word from the main list of distractors

Pseudoword: one of the pseudowords described in the Materials section.

Symbols: a combination of these symbols &#?@$% in random order

Blank: a blank screen (no stimulus) for the same SOA duration.

The response phase began immediately after the presentation of the finishing stimulus. Participants answered two successive questions. First, the Recognition question “Have you seen this word?” was presented in the upper part of the screen along with a probe word presented at the centre of the screen. The probe consisted of the word presented immediately before the finishing item (the mask) in half of the trials and was a new word in the other half. Participants pressed with the right hand the “left arrow key” to indicate YES or the “right arrow key” to indicate NO. Second, the Detection question “Have you seen the target?” was presented and they responded YES or NO using the same keys they used for the Recognition question.

### Results

The first objective of this experiment, as stated above, was to demonstrate that detection performance was disproportionally better than recognition for the short SOA (117ms) relative to the long SOA condition (350ms). To do that we obtained the d-prime values from the Hit and False Alarm rate for both tasks: detection and recognition, see Figure 3. A three-way (Task: detection and recognition, SOA: 117 and 350, Mask: blank, symbols, pseudoword and word) within-subjects ANOVA showed that Task interacted significantly with SOA F(1,29)=10.46, MSE =4.96, p<0.01, η^2^=0.27. Pairwise comparisons showed no effect of SOA in the Detection task (p>0.1), and a significant effect of SOA in the Recognition task, t(1,29)=7.4, p<0.001, d=1.2. Task also interacted significantly with Mask, F(3,87)=14.29, MSE =3.68, p<0.001, η^2^=0.33. In addition, we found that the three main effects reached significance: SOA, F(1,29)=50.26, MSE=14.34, p<0.001, η^2^=0.63; Task, F(1,29)=106.33, MSE=61.64, p<0.001, η^2^=0.79; and Mask, F(1,29)=23.16, MSE =6.83, p<0.001, η^2^=0.44. Finally, the interaction between SOA and Mask approached significance F(3,87)=2.43, MSE=0.64, p=0.07, η^2^=0.08. The three way interaction between Task, SOA and Mask was not significant F(3,87)<1, p>0.1.

**Figure 3:**
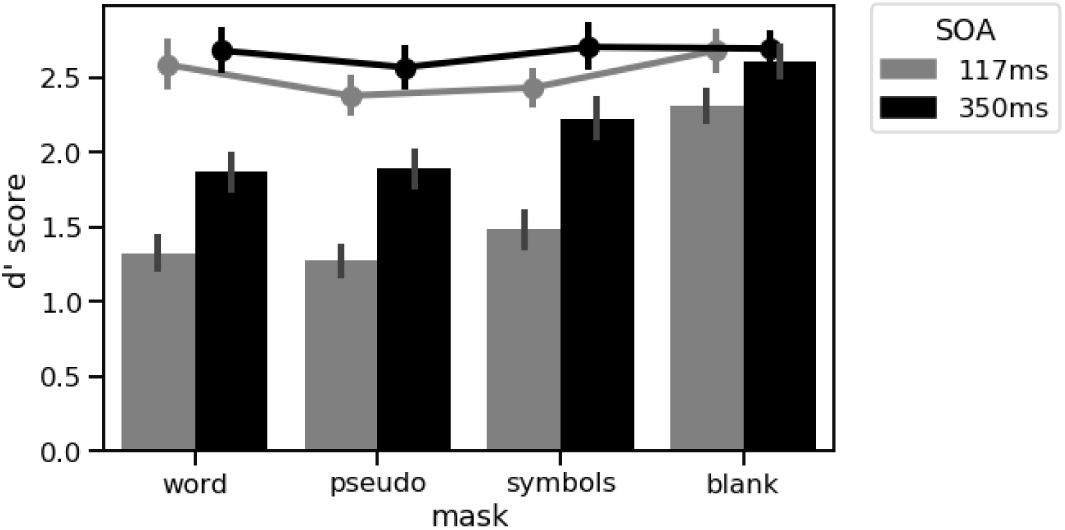
results of experiment 1, comparing recognition (bars) to detection (filled circles) performance across masking conditions and SOA.

The second, more exploratory, objective was to examine to what extend the last stimulus presented in the streams modulated recognition performance in both SOAs. Additionally, to design Experiment 2, we aimed to identify which type of mask would result in higher recognition performance, while still showing the basic interaction between Task and SOA. In this respect, the blank condition (unmasked critical item) was of no interest, since recognition performance was sufficiently high that the interaction between SOA and Task was not significant (p>0.1). Additionally, unmasking the critical item would have made it unrepresentative of the masking of arbitrary distractors in RSVP. For the other masking conditions, the planned pairwise comparisons on the d-prime values of the Recognition task showed that for the 117ms SOA there were no significant differences in all the comparisons among the Words, Symbols and Pseudowords conditions (all ps>0.1). In contrast, in the 350ms SOA condition, the Symbols mask showed larger d-prime values relative to the Words t(29)=2.32, p=0.03, d=0.42 and the comparison between Symbols and Pseudowords approached significance t(29)=1.92, p=0.06, d=0.35.

### Interim summary

As previously discussed, experiment 1 compares detection and recognition performance in a dual task context, i.e. in which participants will be seeking to trade performance on the two tasks off against each other. However, a key question is the memory trace left for an arbitrary distractor in RSVP when participants are solely engaged in detection. Indeed, repeating the recognition question across trials might plausibly have produced a novel demand on participants, namely, for forming a memory trace for each item in the RSVP stream, i.e., not just the target item. Therefore, Experiment 2 considers this question by incorporating a single surprise memory probe on the 14th and last trial (see figure 4), thereby barring the possibility of such an artifact.

**Figure 4:**
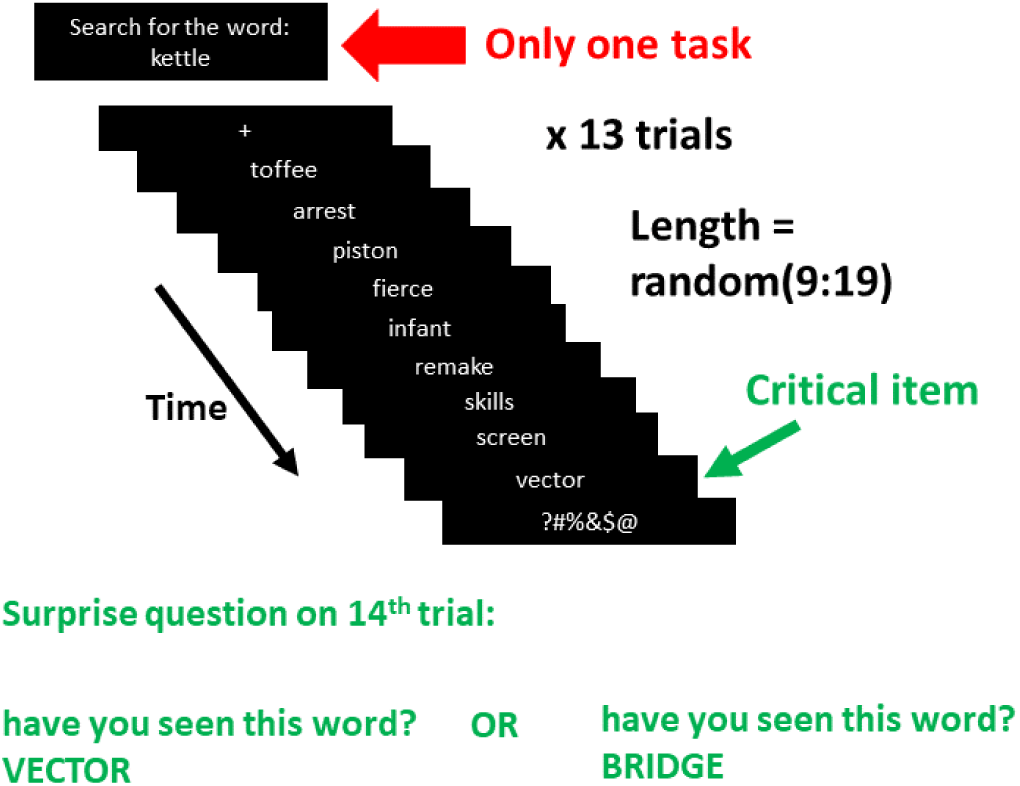
experiment 2 procedure: RSVP streams of words were presented, terminated by a mask. Streams were stopped randomly from the 9th to the 19th item. The task was to detect the specified word and perform a presence/ absence judgement on it at stream end (not shown). Additionally, a surprise question arose on the 14th trial, asking participants to perform a single- probe recognition test on the critical item: the last stream item before the mask.

On the basis of the results of Experiment 1, we selected the symbols mask. The logic here was to use the masking that gives the most capacity to observe recognition memory for distractors. Thus, we selected the mask with the highest memory performance in Experiment 1. If we are able to exhibit low memory performance even for the weakest mask, we will have given the greatest capacity for a memory trace to be observed if it is there.

## Experiment 2

### Methods

#### Participants

151 undergraduate students of the University of Birmingham took part in the experiment in exchange for course credits. 60 (mean age=19.23, sd=1) participants (9 males) took part in the lab session of the experiments. The rest, 91 (mean age=19.65, sd=1.48), took part in the online session. All were native English speakers and had normal or corrected-to-normal vision. The experiment conformed to British Psychological Society criteria for the ethical conduct of research and ethical procedures of the School of Psychology at the University of Birmingham. The experiments performed here were approved by the Science, Technology, Engineering and Mathematics Ethical Review Committee at the University of Birmingham, under ethics programme ERN_20-1291P.

### Materials

Distractors were selected randomly from the main list of distractors of Experiment1. In addition, 30 words were selected for the Recognition test (a list of 15 items to serve as Old words and 15 as New Words). Finally, target words for the Detection task were selected from a list of 30 words.

### Procedure

For the lab version of the experiment, screen model, stimulus presentation software, and stimulus presentation parameters (font and background) were the same as in Experiment 1. The online version was created using PsychoPy (Peirce et al 2019) and hosted by https://pavlovia.org/. The procedure is shown in figure 4. Participants were informed that the experiment consisted of a series of trials, where they would be presented with rapidly presented streams of words. Their task was to search for a specific target word that could be inserted in the streams. At the end of the trial, they were expected to indicate whether they have seen the target.

The experiment consisted of 14 RSVP trials. The structure of each trial was the same as in Experiment 1: Instruction “Search for the word: <TARGET>”, fixation cross, stream of words, (symbols) mask. The SOA of the streams of words was 117ms. The crucial manipulation of this experiment was in the response phase. The first 13 trials ended with the presentation of a Detection question “Have you seen the target word”. As in Experiment 1, participants had to press the “left arrow key” for YES and the “right arrow key” for NO. Crucially, the last trial, 14^th^, was followed by a surprise Recognition question. In this trial, immediately after stimuli presentation, participants were presented with the text “SURPRISE QUESTION, Have you seen this word?” presented at the upper part of the screen along with a probe word presented at the centre of the screen, and the text “press <- to indicate YES or -> to indicate No” presented in the lower part of the screen. The experiment ended after the participants responded to this question.

### Data analysis

Data consisted of the subjects’ responses to the surprise recognition question that ended Experiment 2. 2 subjects were excluded for taking too long to respond (more than 5 seconds). Out of the 58 subjects (and data points) 23 had been presented with a probe corresponding to a New Word and 35 had been presented an Old Word. In addition, out of the 91 subjects who did the online experiment, 42 had been presented with a probe corresponding to a New Word and 49 were presented with an Old Word. We used a model-based approach to estimate Signal-Detection Theory parameters (DeCarlo, 1998), avoiding the need to collapse the data (to get proportions of Hits and False Alarms to obtain the d-prime). In consequence, the responses of the participants were fitted with a generalized linear model (with a Probit link function). The data analysis strategy was designed to test to what extent the data supported the Null Hypothesis of a d-prime value of 0. To do this, as shown in figure 5, we built a Bayesian generalized linear regression model to estimate a posterior distribution of d-prime values. Then, the Bayes Factor was obtained using the Savage-Dickey density ratio (Wetzels et al., 2010) between the prior and posterior estimates.

**Figure 5.**
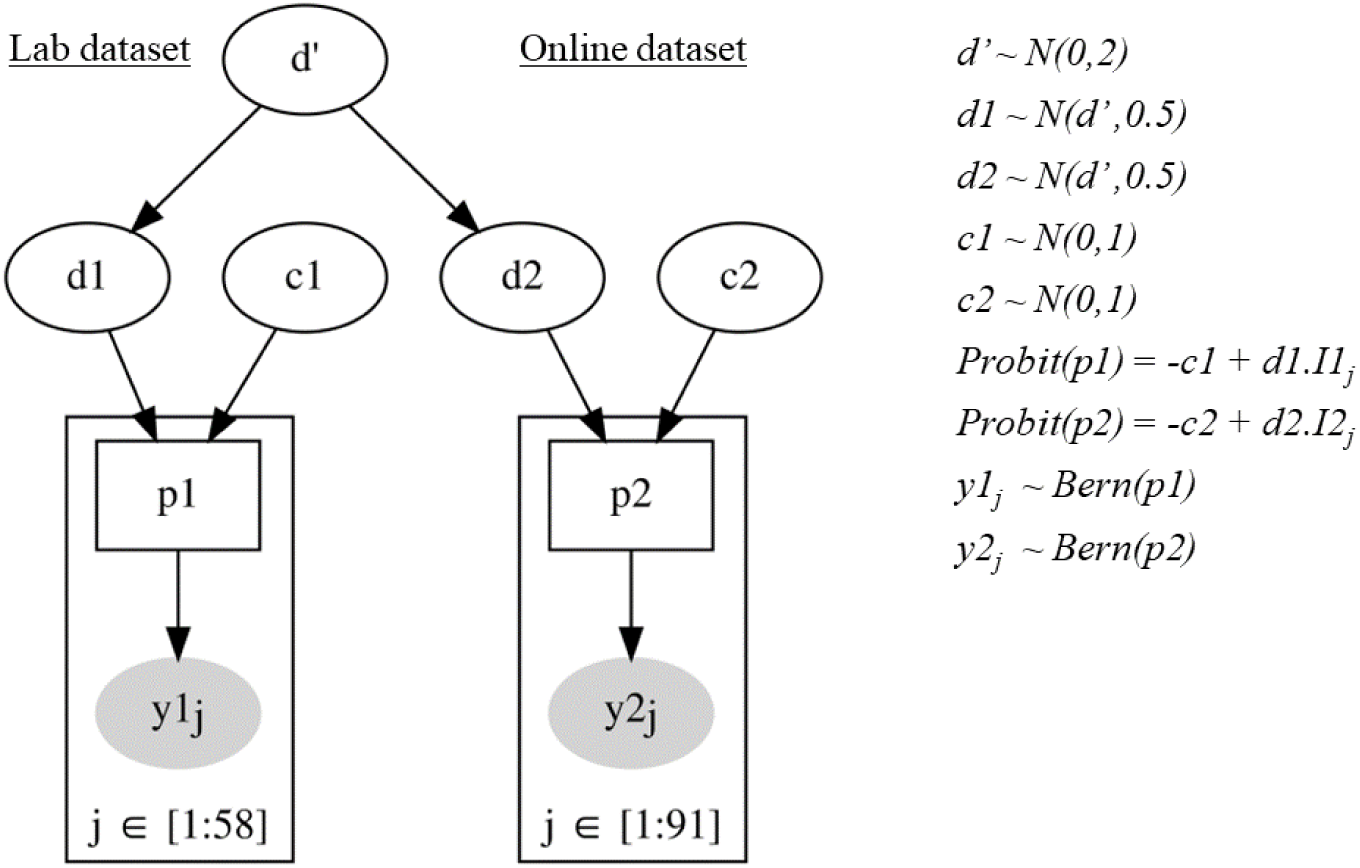
Specification of the Bayesian model for Experiment 2. Index 1 refers to the in-person (Lab) experiment and index 2 the online version. For each participant (j), y1_j_ and y2_j_ are either a one or a zero, depending upon whether the jth participant responded with YES or NO on their 14th trial. p1 and p2 are probabilities of a YES response across all 14th trials. d’ is the highest-level d- prime, which is a prior for d1 and d2, the d-primes for each of the two datasets (in-person and online). c1 and c2 are the criteria for the two datasets, and I1j and I2j indicate target presence/absence on 14^th^ trial of participant j. The negative of each criterion gives the intercept, and the relevant d-prime, the slope of the probit regression. *Bern(pi)* indicates a Bernoulli distribution, for which 1 has a probability of pi and 0 a probability of (1-pi).

The second objective of the analysis was to test whether the results of Experiment 2 were significantly different from those of Experiment 1. To do this, we selected the data from Experiment 1 where the same SOA and Mask conditions as in Experiment 2 were used. To fit the data from Experiments 1 and 2, we extended the Bayesian model from Experiment 2. The part of the model that fits the data from Experiment 1 is a Hierarchical version of the model for Experiment 2, where each participants’ parameters are drawn from parent distributions. d-prime values were estimated for both experiments. The Bayes Factor was then obtained by using the Savage-Dickey density ratio between the Effect Size density estimated from the priors and from the posterior at 0.

### Results

#### Evidence for the Null Hypothesis (d-prime=0)

We first focussed on the Experiment 1 Bayesian model presented in Figure 5. For the lab version, the proportion of Hits was 0.57 and the proportion of False Alarms was 0.43 (d-prime: 0.34). For the online sample, the proportion of Hits was 0.61 and the proportion of False Alarms was 0.45 (d-prime = 0.40). To estimate the posterior of the parameters, 5000 draws (tuning samples = 300) were sampled from it using the No U-Turns Sampler (NUTS) (Hoffman & Gelman, 2014). With a normal distribution (mean=0, sd=2) as a prior for the d-prime, a Bayes Factor of 3.29 (see Figure 7) was obtained in favour of the Null Hypothesis and the mean of the posterior of the d- prime was 0.36 (94% credible interval = -0.41 to 1.12). In addition, to assess the robustness of the Bayes Factor, we fitted the model using a Uniform prior (lower bound=-3, upper bound=3). In this case, the Bayes Factor in favour of the Null Hypothesis (d-prime=0) was 3.84. The mean of the posterior of the d-prime was 0.38 (94% credible interval = -0.38 to 1.17).

#### Difference between Experiments 1 and 2

The standardized effect size of the difference between the d-prime estimates of Experiments 1 and 2 was obtained by sampling 3000 draws (tuning=300 draws) from the posterior of the model depicted in Figure 6. Similarly, 3000 draws were obtained to estimate the Effect Size from the priors of the model. The Bayes Factor was the ratio at 0 (no difference between Experiments) between the estimates of the Effect Size from the posterior and priors. The Bayes Factor was 12.87 (see Figure 8), which indicates strong evidence for the Alternative Hypothesis of a significant difference between the performances of the two experiments.

**Figure 6.**
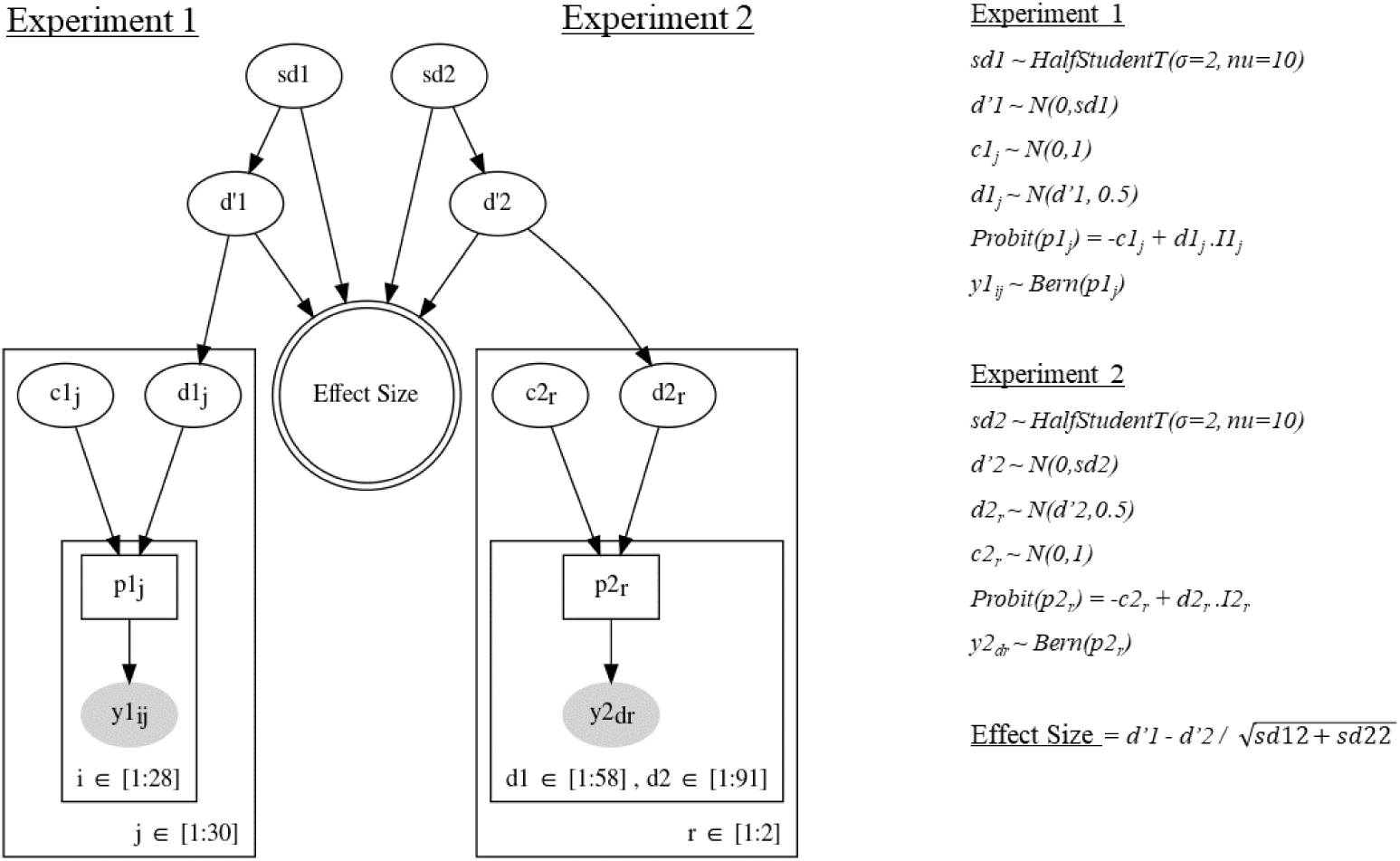
Bayesian hierarchical model for Experiments 1 and 2. Index 1 refers to Experiment 1 and index 2 to Experiment 2. In Experiment 1, for each participant (j) and trial (i), y1_ij_ is either a one or a zero, depending upon whether the participant responded with YES or NO on the recognition question. In Experiment 2, for each version of the experiment (r, in-person and online) and each participant (d, which corresponds to d1 in first version of experiment and d2 in second version), y2_dr_ is either a one or a zero, depending upon whether the participant responded with YES or NO on the 14^th^ trial. d’1 and d’2 are the highest-level d-primes from both experiments, which are priors for d1_j_ and d2_r_ respectively, the d-primes for each participant in Experiment 1 and for each dataset (in-person and online) in Experiment 2. c1_j_ is the criterion for each participant of Experiment 1, and c2_r_ is the criterion for each dataset of Experiment 2. I1_j_ and I2_r_ indicate target presence/absence. The negative of each criterion gives the intercept, and the relevant d-prime, the slope of the probit regression. *Bern(p_j_)* and *Bern(p_r_)* indicates a Bernoulli distribution, for which 1 has a probability of p_j_ (or p_r_, for *Bern(p_r_)*) and 0 a probability of (1-p_j_) or (1-p_r,_ for *Bern(p_r_)*).

**Figure 7.**
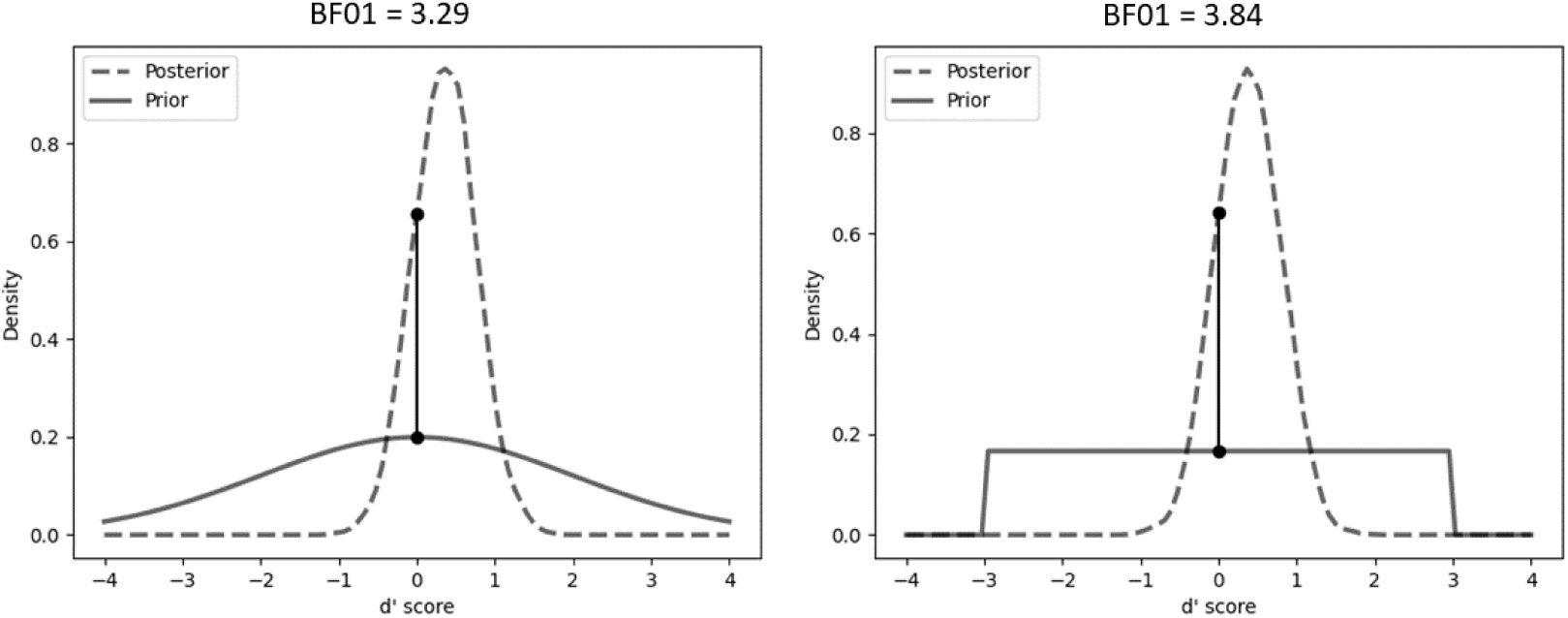
Posterior (dashed line), prior (solid) and Bayes Factor (density ratio, at vertical line at 0), Experiment 2. Left, Normal Prior (mean=0, std=2). Right, Uniform Prior (-3,+3).

**Figure 8.**
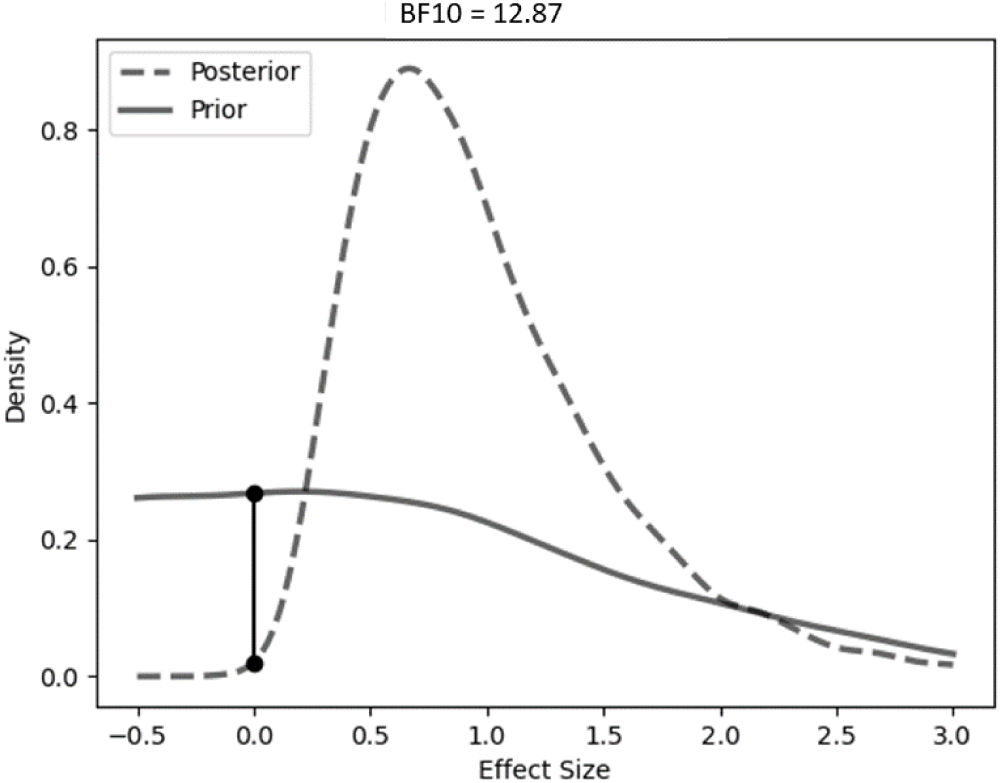
Posterior distribution (dashed line), prior distribution (solid) and Bayes Factor (density ratio, at vertical line at 0) of the Effect Size of the comparison between Experiment 1 and 2.

## Discussion

### Summary of findings

In these two experiments, we compared detection (for targets) and recognition (of distractors) performance in RSVP. An important finding was an interaction between Task and SOA in Experiment 1, whereby the change in SOA did not impact Detection, but substantially impacted Recognition. This replicates the finding of Bowman and Avilés (2021), where it was also argued that the interaction is unlikely to be driven by a ceiling effect on Detection performance. This interaction sits well with our perspective that Detection performance is high when the brain is searching subliminally for targets (SOA 117) and not much higher, if at all, when searching supraliminally (SOA 350), indeed discrimination (d’) does not change between these two SOAs (Avilés et al., 2020). However, recognition improves dramatically from SOA 117 to SOA 350, suggesting a regime change for recognition between subliminal and supraliminal. This fits with our theoretical perspective that the brain can search subliminally for what it “knows”, this is subliminal salience search, but consciousness is required when laying down long lasting memory traces.

It is also the case that as expected, d-prime values were significantly lower for Recognition compared to Detection (see Figure 3) independently of the SOA or the type of mask that followed the critical items. Furthermore, recognition performance was clearly above chance in Experiment 1 even for the shortest SOA. However, in Experiment 2, we found evidence that recognition performance is extremely low (see Figure 7, for the posterior estimates of the model) when the recognition question was not expected by the participants. That is, in the context of a single task (Detection), recognition performance was on the basis of a Bayesian test, not different to chance, and significantly lower than the d-prime values obtained when Recognition was part of the task-set, as in Experiment 1.

### Comparing recognition and detection

Central to this paper is our comparison of d-primes for detection and for recognition. How can we conceptually justify this comparison, since these are two different cognitive functions? Essentially, d-prime in RSVP, indexes the capacity to discriminate the presence of a critical item appearing at a single arbitrary position in a stream (quantified as the Hit rate) versus not appearing at any position in the stream (quantified as a Correct Reject). The difference between the two is that, for detection, the critical item is the target being searched for, while, for recognition, it is the (single) item probed in the recognition memory display. In addition, although our recognition test probes the last item (before the mask) in the RSVP stream, our approach of randomly stopping the stream, to prevent any capacity to predict when the end is coming, simulates (in the mind of the participant) the occurrence of the probed item at any arbitrary position in the stream.

Thus, essentially, the detection and recognition d-primes are assessing the same discrimination, with the only difference being whether the critical item is the (detection) target or the item "post probed” (i.e., penultimate in the stream).

### Relationship to our Previous Work

Bowman and Avilés (2021) was the first article to extended Potter (1976)’s findings by firstly, probing memory specifically for items at the end of the stream, in order to reduce the possibility for forgetting, and secondly, by varying the length of RSVP streams, in order to prevent participants from predicting their ending. Bowman and Avilés (2021) focussed on characterising recency and comparing it between recall and recognition. A striking finding was that at RSVP rates, recall was, firstly, extremely poor and secondly, exhibited a close to maximal recency effect, in the sense that the little that was recalled was almost exclusively at position -1, i.e. almost no items at positions - 2 and -3 were recalled.

Most importantly, though, the poor recall performance identified in Bowman and Avilés (2021), suggests that the brain’s capacity to detect and identify targets in RSVP, with detection performance often around 80%, cannot be justified by a recallable level of memory imprint. In this article, the -1 item could be recalled on less than one in five streams (i.e. <20%).

This excludes free recall as a source of a memory imprint left by distractors during the process of determining that they are not targets. However, recognition performance was higher in Bowman and Avilés (2021), leaving the possibility that rejected distractors leave a familiarity, rather than recollection trace, an idea that would be consistent with the glance phase of the Glance-Look model of RSVP perception Su et al (2011). This possibility that there is a familiarity trace is what we have specifically focused on in this paper.

Additionally, we refined the experimental paradigm employed in Bowman and Avilés (2021). Firstly, we only probed recognition memory, recall was not considered. Secondly, we probed memory with a single item at the end of each RSVP stream. This is to avoid the disruption that can arise in multiple-item probes, where each memory retrieval can potentially disrupt the memory for other items being probed; see, for example, the effect of retro cues in iconic memory experiments (Sligte et al, 2008; Sperling, 1960). Thirdly, we only probed the -1 item (i.e. the last before the mask) in the RSVP stream, as a result of which we have not considered recency in this paper. Finally, we explored different types of end of stream mask.

In pilot work, we determined that the mask used in Bowman and Avilés (2021), a sequence of hashes, gave substantially higher performance than variable-character masks. However, the central idea of our experiments is that the distractor that we probe should be presented with a similar perceptual difficulty to an arbitrary distractor in the stream, which would be backward masked by another distractor. Accordingly, here we preferred an end of stream item that is a variable-character mask. We explored different possible letter masks in experiment 1 and selected the symbols mask, since it has the highest performance across the variable-character masks, enabling us to give as much chance of high performance being observed as possible.

### Surprise Experiment

Our second experiment seeks to assess the memory left for an arbitrary RSVP item, while the brain is in its “native” subliminal search state, i.e. it is just performing a detection task. Although we have presented evidence for zero memory for RSVP distractors (Bayes Factor in favour of d-prime=0 was 3.84), we do acknowledge the need for a replication of this effect. Indeed, it seems surprising that there really is a complete absence of a memory imprint. However, what we believe we can argue is that the memory imprint is at least weak and is certainly very substantially smaller than detection performance at RSVP rates. This is sufficient for the line of argument we are making here, i.e. that the brain can exclude distractors in RSVP with very little in the way of a memory trace being left for those distractors.

An alternative explanation of the poor performance in the surprise recognition question is that this is a simple case of forgetting. The retention interval (the gap between presentation and probe) and the presence of the surprise question might be sufficient to disrupt the retrieval of the critical item. More evidence is required to reject this hypothesis, but initial evidence seems to rest credibility from the forgetting explanation. Chen and Wyble (2016), using a similar surprise question method, showed successful retrieval of perceptual attributes that participants did not expect to report. That is, the surprise question did not disrupt the memories for attributes that needed to be encoded for other purposes than report. They concluded that the poor performance in a surprise recognition question demonstrates a failure of memory consolidation rather than forgetting.

Furthermore, we believe that the brevity of the retention interval might be sufficient to reject the forgetting hypothesis. Indeed, evidence from RSVP Event Related Potential (ERP) experiments suggest latencies for conscious perception. In RSVP, the ERP component most commonly associated with conscious perception is the P3b (Bowman et al, 2013, 2014; Craston et al, 2009; Vogel et al, 1998; Sergent et al, 2005); see, for example, Pincham et al (2016) and Jones et al (2020) for particularly direct support for this association. Across the numerous RSVP ERP experiments, the P3b is rarely if ever, found to start before 350ms and to peak before 450ms after the eliciting stimulus.

If we extrapolate these P3b latencies to our experiments, we are left with the scenario that conscious perception of the -1 item would occur perhaps 170ms *after* the memory probe started being presented^2^. In this context, is the forgetting hypothesis still tenable? It would require the -1 stimulus to be perceived and forgotten all while the memory probe screen was being displayed, and indeed had already been presented for some 170ms before the perception-forgetting would start unfolding.

### Late Selection

Our theoretical position arises from the Simultaneous Type/ Serial Token (STST) model (Bowman & Wyble, 2007; Bowman, Wyble, Chennu & Craston, 2008). The findings reported here are consistent with the STST model, while also resonating with a long tradition in attention research, so called *late attentional selection* theories (Deutsch & Deutsch, 1963; Duncan, 1980). However, the role of conscious experience is more overtly emphasized here, than in the classic late *attentional* selection theories.

Duncan (1980) is a key demonstration of late selection. A first parallel stage (which would correspond to stage-1, extracting types, in STST) is proposed to precede a capacity limited second stage (which would correspond to the serial second stage in STST). However, late selection theories argue that as well as simple stimulus properties, such as colour, the first system can extract form and meaning. This enables a full identification of a stimulus, exactly what is required in a subliminal search system, that identifies salient stimuli pre-consciously.

Evidence for late selection is typically taken from (pop-out) visual search (Wolfe & Horowitz, 2017) and divided attention experiments, as studied for example, with dichotic listening (Broadbent, 1956). Indicative findings in dichotic listing are that effects of the non-attended stream are substantial, e.g. cocktail party effect (Moray 1959).

A highly relevant finding is that stimuli can be rejected as non-targets/ non-salient in parallel, but multiple stimuli cannot be *selected* in parallel, because the former can be performed by the first (parallel) system, but the latter requires the second (limited-capacity) system. Consistent with this, performance is substantially better with a concurrent correct rejection than with a concurrent hit (Duncan, 1980). Thus, the interference cost of rejecting as a non-target is minimal, or, in other words, one can reject stimuli in parallel, but one cannot "detect" multiple stimuli in parallel!

Our finding that memory traces for (rejected) distractors are low relative to detection performance is wholly consistent with this aspect of late selection, where presumably, a distractor can be easily rejected without it leaving a strong memory trace. This distinction between the processing of distractors and targets naturally suggests that in RSVP, rejection of distractors can be performed subliminally, while once identified/detected, targets are consciously perceived. This is what we call subliminal salience search (Bowman et al, 2013).

However, STST adds to these late selection theories, by providing an explicit explanation of why the second stage is limited in capacity. In STST, the need to unambiguously associate salient stimuli to episodic information is what induces the capacity limitations. In more general terms, in order that memories of our experiences are sequenced, our experiences themselves need to be sequenced; this is the serial token allocation of the second stage of STST. We elaborate on this aspect of STST and the resulting tokenized-percept hypothesis next.

### Conclusions

The findings presented here and in Avilés et al. (2020), Bowman and Avilés (2022) and Bowman and Avilés (2021), suggest a marked fragility (indeed potentially absence) of memory for incidental stimuli presented in RSVP. It is clear that *salient* stimuli in RSVP streams frequently create durable traces, enabling accurate presence/absence or identification judgements on targets. However, on the basis of our work, it seems that the brain can propel stimuli of interest into consciousness and working memory, without incurring a memory storage “cost” for stimuli that are not of interest. In this sense, the brain successfully applies a strong salience filter (Lachter, Forster, & Ruthruff, 2004; Bowman & Wyble, 2007).

We have talked in terms of memory for incidental stimuli being a “cost”. This is true if one views the task of the perceptual system as to act as a filter; however, the quotation marks here are pertinent, since this failure in incidental encoding does rule out one valuable functional capacity, and that is, the capacity to attribute episodic information to stimuli. As an illustration, let’s take the ability to know that a stimulus has been previously presented (i.e. that it is a repetition) as a classic example of an episodic task. A repetition task can only be effectively performed if there is *incidental durability*. That is, the stimulus only becomes salient at the point of its repetition; however, to know that it has been repeated, a previous instance of occurrence has to have left a memory trace, and at the point of that first occurrence it was *not* salient. Does this indicate a key duality between the subconscious and the conscious: the subconscious can find stimuli in our environments with exquisite accuracy, while only the conscious can exhibit incidental durability enabling episodic information to be represented? The former of these is subliminal salience search, and the latter what we call the *tokenized percept hypothesis* (Avilés et al., 2020), Bowman and Avilés (2021), which has similarities to a hypothesis that Kanwisher has presented (Kanwisher, 2001). At its heart, the tokenized percept hypothesis claims that a central function of conscious experience is to episodically tag experiences in time and indeed space, and that capacity is uniquely conscious.

## Footnotes

1 An SOA of 117ms is in the typical range in the RSVP paradigm; see Potter, 1976 or Cecotti et al., 2014 for reviews

2 The latency of the P3b suggests an approximate time to conscious perception of 400ms from the -1 item onset. The memory probe onsets 230ms after the onset of the -1 item. 400 minus 230, gives us 170ms.

## Notes

### Competing Interest Statement

The authors have declared no competing interest.

